# SC-BIG: A Hierarchical Bayesian Model for Bulk-Informed Single Nucleotide Variant Calling in Single Cells

**DOI:** 10.64898/2026.03.12.705671

**Authors:** Daniel Schütte, Thomas J. Y. Kono, Roland F. Schwarz

## Abstract

Single-cell DNA sequencing (scDNA-seq) has emerged as a primary method for studying the evolution of cancer genomes and intra-tumor heterogeneity. However, despite technological advances, scDNA-seq remains noisy and is affected by amplification biases and allelic dropouts. Accurately determining the presence or absence of candidate somatic nucleotide variants (SNVs) in individual cancer cells therefore remains challenging. One strategy to alleviate this issue is to perform bulk whole-genome sequencing simultaneously with single-cell sequencing. To date, only few computational methods have been developed for bulk-informed detection of somatic SNVs in single cells, and existing methods do not adequately account for somatic copy-number alterations or clonal admixtures.

We here present SC-BIG, a hierarchical Bayesian model that leverages bulk sequencing data from a representative tumor sample to improve SNV detection. SC-BIG propagates uncertainty across multiple biological parameters, including copy number alterations, sample purity, and SNV clonality. In a first step, the cancer cell fraction (CCF) of a SNV is jointly estimated from bulk and single-cell data. The CCF in turn then acts as a prior in the second inference step to calculate per-cell posterior probabilities for the presence of the SNV. We demonstrate that across simulated scenarios of varying CCFs, SC-BIG outperforms both naive thresholding and ProSolo, the only bulk-informed single-cell mutation caller described so far. Importantly, SC-BIG produces well-calibrated posterior probabilities that provide interpretable uncertainty quantification, enabling direct integration into downstream analyses such as phylogenetic reconstruction.

## 1 Introduction

Single-cell DNA sequencing (scDNA-seq) of cancer samples has enabled the study of intratumor heterogeneity and clonal dynamics at very fine resolution and scale [1]. This is accompanied by increased noise from sparse coverage, amplification errors, and allelic dropout (ADO). Existing bioinformatics methods address these challenges through various strategies. *Monovar* [2] discovers SNVs *de novo* using a Bayesian model that incorporates ADO and aggregates evidence across multiple cells. *SCcaller* [3] and *SCAN-SNV* [4] estimate local amplification bias from germline heterozygous SNPs (hSNPs) before applying bias-corrected variant calling.

While these methods enable *de novo* SNV discovery from isolated single-cell experiments, tumor sequencing increasingly comprises multiple data modalities, including contemporaneously sampled scDNA-seq and bulk DNA-seq [5–8]. Lähnemann *et al*. [9] were the first to attempt bulk-informed variant calling with *ProSolo*, which probabilistically genotypes variants in single-cell data using bulk data as a prior. ProSolo builds on the Bayesian framework Varlociraptor [10] and partitions the joint single-cell and bulk genotype space into seven mutually exclusive events. The model assumes one of three diploid allele frequencies (***θ***_***s***_ ∈ **{0, 0.5, 1}**) for each cell and a discretized bulk allele frequency ***θ***_***b***_ ∈ **[0, 1]**. Events correspond to the three diploid genotypes, dropout of either allele, and sequencing errors of either allele. To account for whole-genome amplification artifacts in the single cells, ProSolo’s error model is based on the coverage-dependent beta-binomial model of Lodato *et al*. [11]. This model was fitted to empirical multiple displacement amplification (MDA) data and captures allelic imbalance through a mixture of beta-binomial distributions whose shape parameters are linear functions of read depth.

Despite its sophistication, ProSolo makes two strong assumptions that are frequently violated in cancer genomics data. First, all sites are assumed to be diploid: the Lodato model restricts the true single-cell allele frequency to ***θ***_***s***_ ∈ **{0, 0.5, 1}**, making it unable to represent non-diploid copy number states. Second, each variant in the bulk sample is assumed to be explainable by a mixture of at most 2 genotypes that differ by a single mutational step. These assumptions can lead to miscalibrated posteriors for subclonal variants at loci with copy number alterations.

Here, we develop SC-BIG, a method to leverage complementary bulk sequencing data to estimate variant presence in individual single cells. Given bulk tumor sequencing, scDNA-seq data, and optionally a set of hSNPs, we compute the probability that a specific candidate mutation observed in bulk is present in a single cell. SC-BIG relaxes both of ProSolo’s key assumptions: it supports copy number states ***C*** ∈ **{1, …**, **10}** with arbitrary multiplicity ***m* ≤ *C***, and it explicitly models subclonal variants by marginalizing over cancer cell fraction. Because CCF suffers from inherent undecidability, a prior is estimated from a pseudo-bulk of cells with high coverage. To simplify the inference algorithm, we assume that all cells are malignant and focus on variant presence rather than estimating an exact variant allele frequency (VAF). We use simulated data to provide a proof of principle and show that SC-BIG performs favorably compared to both naive VAF thresholding and the ProSolo implementation of Lähnemann *et al*. [9].

## 2 Methods

SC-BIG infers the probability that a variant of interest is present in a specific single cell, given bulk evidence. For each variant, it takes as input bulk and single-cell read counts, an estimate of local copy number ***CN*** in the bulk, tumor purity (***π***), and sequencing error ***ϵ***. It optionally takes read counts at linked germline heterozygous SNPs (hSNPs; see Table 2, Figure 7 in the Appendix). Inference of variant presence then proceeds in three steps (Figure 1). First, an empirical prior on cancer cell fraction (CCF) is constructed from the aggregate fraction of cells showing variant reads. Second, a CCF posterior is obtained from bulk data by marginalizing over ***CN***, multiplicity ***m***, and ***π*** via a Markov Chain Monte Carlo (MCMC) procedure. Third, per-cell variant probabilities are computed by combining the CCF posterior with a single-cell likelihood that accounts for amplification bias under an error model. SC-BIG implements its own simple, technology-agnostic beta-binomial model for which it uses hSNPs as well as the MDA-specific model of Lodato *et al*. [11].

**Figure 1:**
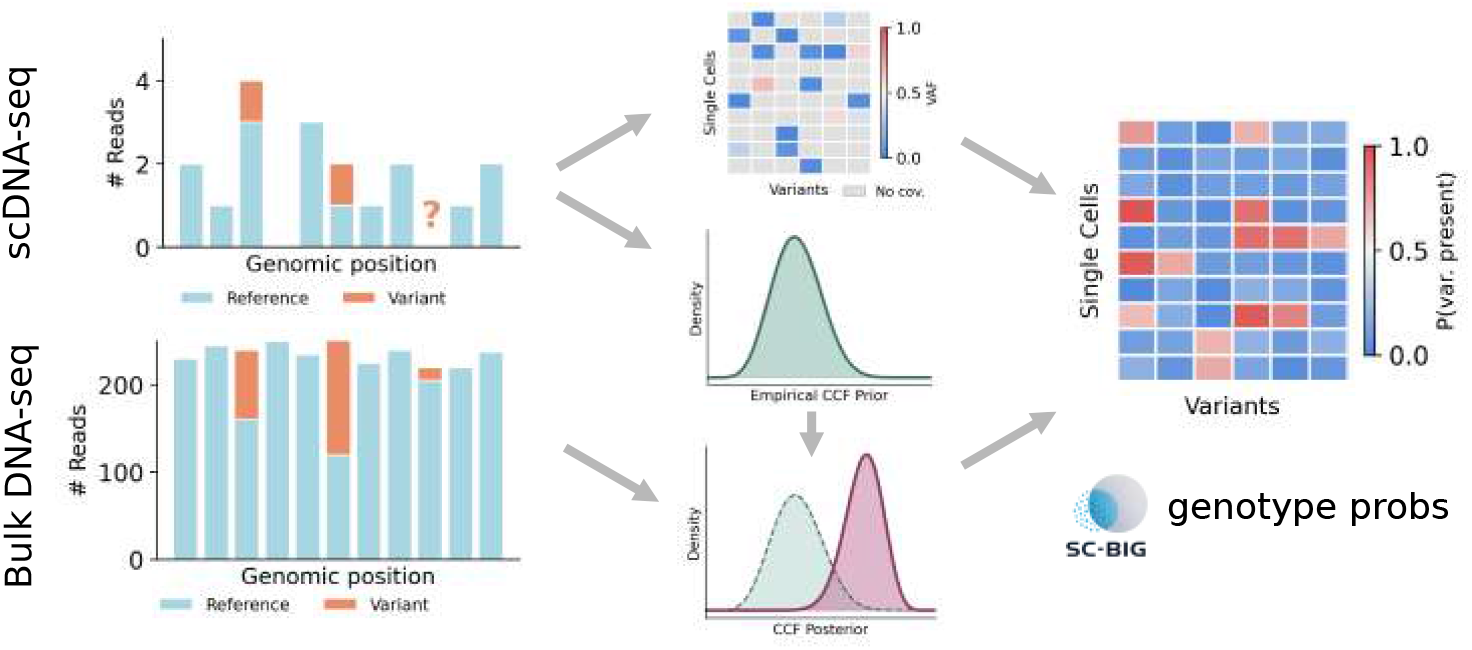
SC-BIG method overview. Starting from sparse single-cell reads at a candidate variant site, SC-BIG constructs an empirical CCF prior from single-cell aggregates and a VAF matrix across cells and variants. Bulk read counts are combined with the CCF prior to obtain a CCF posterior via MCMC. Then, per-cell variant probabilities are computed based on observed reads, an error model, and the CCF posterior.

### 2.1 Model Formulation

We will now describe the SC-BIG model in more detail. Figure 2 provides the corresponding Bayesian network diagram. Let ***z*** ∈ **{0, 1}** indicate variant presence in a single cell:

**Figure 2:**
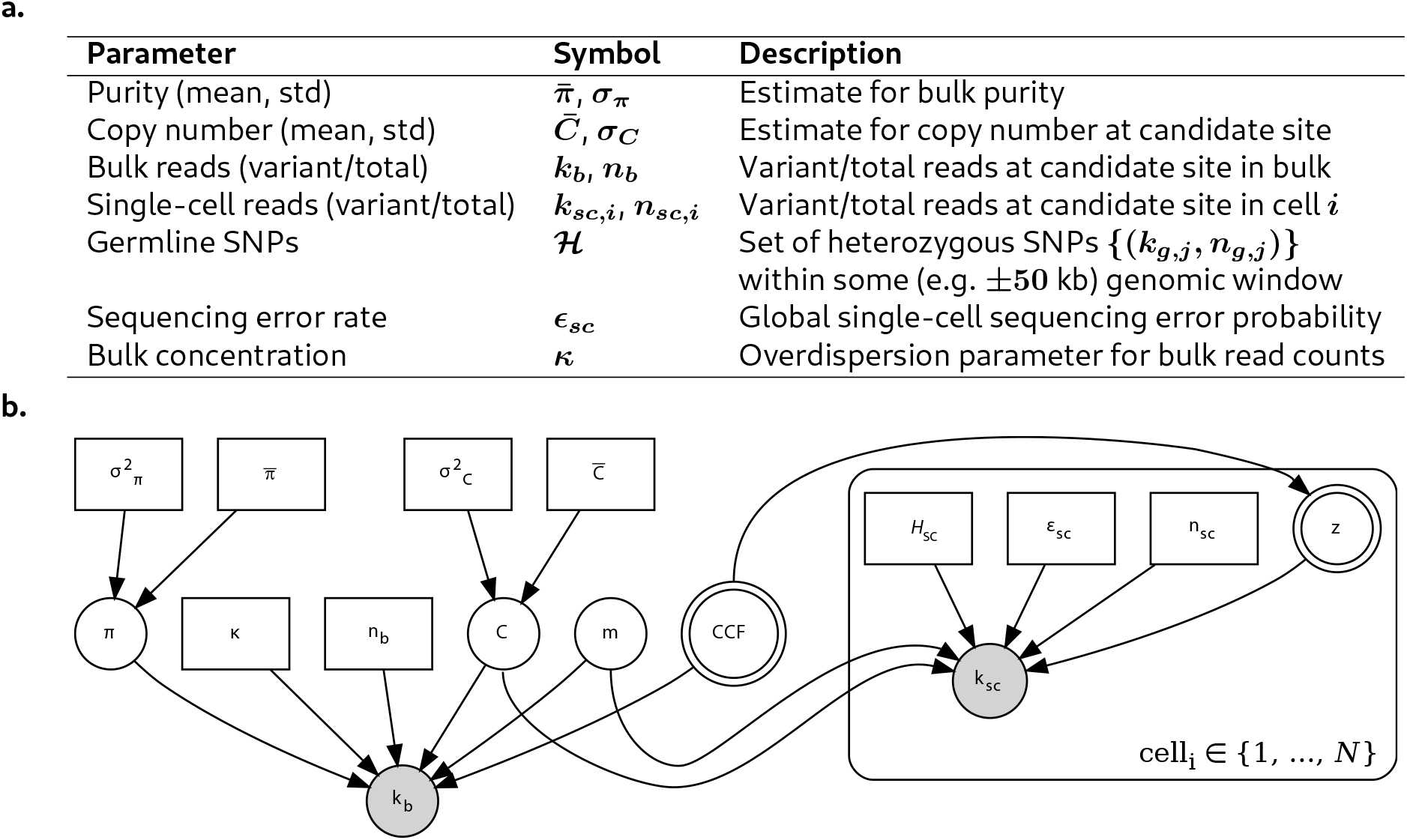
Model overview. (a) Required input data for SC-BIG. (b) Bayesian network diagram. Double circles indicate primary inference targets. Shaded and unshaded circles represent observed and latent random variables, respectively. Rectangles represent fixed input data.

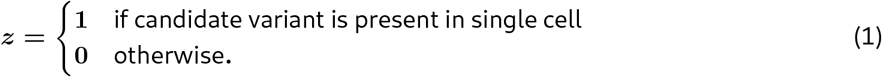

We assume that the posterior probability of variant presence depends on both single-cell and bulk data

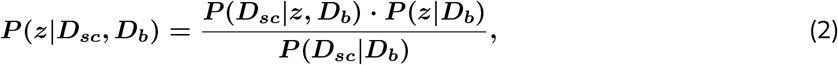

Let us first consider the case of ***z* = 1**. Under SC-BIG’s model, ***D***_***sc***_ comprises the number of total and variant reads and bulk evidence ***D***_***b***_ is quantified via the latent cancer cell fraction. The latter we thus marginalize over:

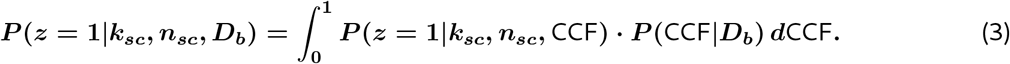

To calculate these terms, we make the key assumption that ***P* (*z* = 1** | CCF**) =** CCF: a malignant cell harbors the mutation with a probability equal to the variant’s cellular prevalence in the bulk sample. Marginalizing over ***P* (**CCF|***D***_***b***_) in equation (3) then propagates uncertainty from bulk to single-cell inference. For a given CCF value, we use Bayes’ theorem once more to compute ***P* (*z* = 1**|***k***_***sc***_, ***n***_***sc***_, CCF) as

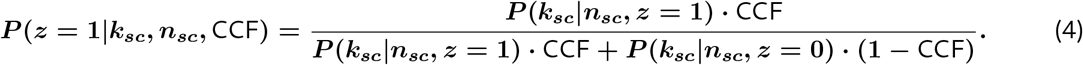

Next, the single-cell likelihood terms ***P* (*k***_***sc***_ | ***n***_***sc***_, ***z*** ∈ **0, 1**) in equation (4) are defined in either one of two ways. We will start with the simpler error model implemented by SC-BIG to facilitate simulation and increase applicability across single-cell sequencing technologies:

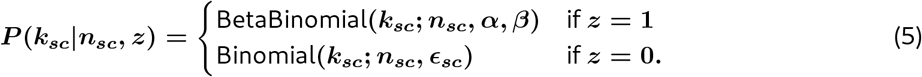

The beta-binomial likelihood accounts for amplification bias, because whole-genome amplification introduces stochastic deviations from binomial expectations [3, 4]. We estimate distribution parameters from germline heterozygous SNPs (hSNPs), assuming a 1:1 allelic ratio. More specifically, beta-binomial parameters ***α*** and ***β*** are decomposed into the concentration parameter ***τ* = *α* + *β*** that quantifies strength of amplification bias and an expected VAF ***α/*(*α* + *β*) = *m/C*** determined by copy number ***C*** and multiplicity ***m***. This parameterization preserves the biologically expected mean VAF while capturing overdispersion. To identify ***τ***, our estimation procedure uses pooled hSNPs across all available single cells and fits a symmetric beta-binomial (***α* = *β* = *τ/*2**) via maximum likelihood estimation. Given the resulting ***τ***, we parameterize the somatic variant beta-binomial as:

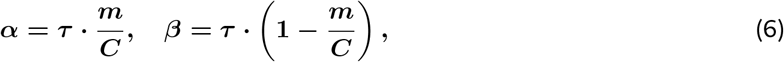

ensuring mean VAF = ***m/C*** while ***τ*** captures population-level overdispersion.

Importantly, a single beta-binomial might not fully capture amplification bias and parameter estimation might not be feasible for real data. Thus, SC-BIG supports the coverage-dependent amplification bias model of Lodato *et al*. [11] as a well established alternative for MDA data. It uses a mixture of beta-binomial distributions with shape parameters that are linear functions of read depth. Because the Lodato model is defined only for diploid allele frequencies ***θ***_***s***_ ∈ **{0, 0.5, 1}**, we map the true allele frequency ***m/C*** to the nearest supported value:

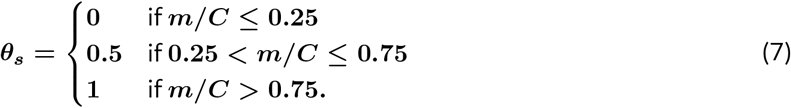

For a given ***θ***_***s***_, SC-BIG follows ProSolo’s model by specifying a distribution over a distorted allele frequency ***ρ***_***s***_. Per-read likelihood is evaluated at ***ρ***_***s***_ using a binomial model with base error rate ***e*** and combined with Lodato’s coverage-dependent beta-binomial likelihood

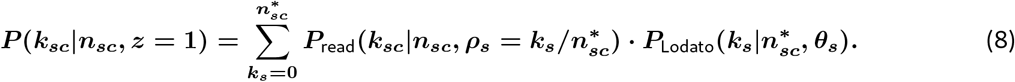

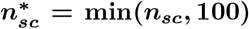, because the Lodato model caps coverage at ***n***_***sc***_ **= 100**, and ***P***_Lodato_ is the Lodato beta-binomial probability mass function. ***P***_read_ is the per-read likelihood defined by ProSolo:

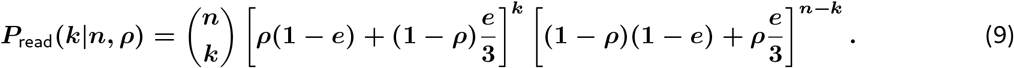

All parameters are taken from ProSolo’s source code. When the Lodato model is used, the variant-absent likelihood ***P* (*k***_***sc***_ | ***n***_***sc***_, ***z* = 0)** is also computed via equation (8) with ***θ***_***s***_ **= 0**, rather than the simple binomial of equation (5), ensuring that both ***z* = 1** and ***z* = 0** likelihoods account for amplification artifacts consistently. This error model is well-characterized for MDA-derived data and enables direct comparison with ProSolo, because SC-BIG can use it during both inference and simulation. The beta-binomial model of equation (6), which makes no technology-specific assumptions about amplification errors, serves as a fall-back for other scDNA-seq technologies, such as primary template-directed amplification **??**.

To compute equation (3), we need the CCF posterior ***P* (**CCF|***D***_***b***_). Bulk sequencing provides evidence about population-level CCF, but uncertainty in variant multiplicity ***m*** potentially leads to a multimodal posterior. For example, a bulk VAF of 0.4 is consistent with both ***m* = 1**, CCF **≈ 0.8** (heterozygous variant in 80% of cells) and ***m* = 2**, CCF **≈ 0.4** (homozygous variant in 40% of cells). This is because the expected bulk variant allele frequency depends on ***π, C, m***, and CCF as follows [12]:

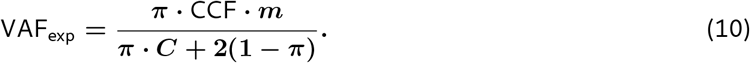

On the other hand, the likelihood of observing ***k***_***b***_ bulk variant reads is computed as:

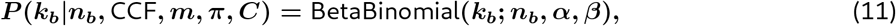

with ***α* = *κ* ·** VAF_exp_, ***β* = *κ* · (1 –** VAF_exp_), and ***κ*** being a concentration parameter for bulk overdispersion. Consequently, CCF posterior estimation will have a major impact on inference accuracy in SC-BIG. While the CCF prior ***P* (**CCF) can be set to uniform, we can also use aggregated observations of the single cells to obtain a more informative, empirical prior. The fraction of single cells showing variant reads provides evidence about population-level CCF independent of multiplicity. Thus, we implement the following Beta distribution as an alternative to a uniform CCF prior:

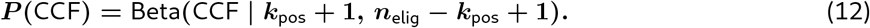

***k***_pos_ is the number of cells with at least one variant read (***k***_***sc***_ **≥ 1**) and ***n***_elig_ is the total number of cells with sufficient coverage (***n***_***sc***_ **≥ 3**). This formulation provides natural regularization because with little reliable single-cell data, the prior converges to Beta**(1, 1)**, which is equivalent to a uniform prior over **[0, 1]**. Given either one of the CCF priors, the CCF posterior is obtained by marginalizing over ***π, C***, and ***m***:

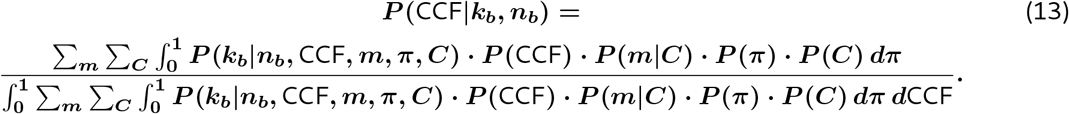

The prior distributions for ***C, π***, and ***m*** are specified as follows:

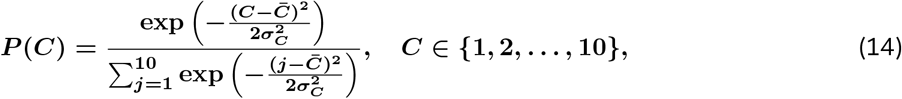

and

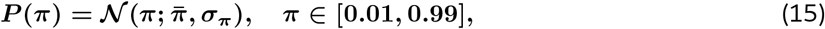

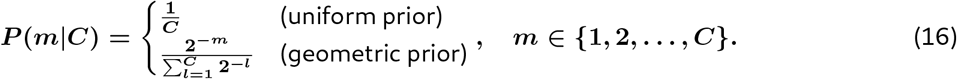

The geometric prior ***P* (*m*** | ***C*) ∝ 2**^−***m***^ favors lower multiplicities, reflecting the expectation that most somatic variants are heterozygous (***m* = 1**), and is used for all inference runs presented in this manuscript. Algorithm 1 in the Appendix summarizes the complete inference procedure.

### 2.2 Simulation Design

Algorithm 2 in the Appendix is used to generate synthetic data for validation. To produce a realistic distribution of cellular prevalences, we use the Kingman coalescent [13] to simulate a genealogy of tumor cells. Starting from ***N*** leaf nodes, pairs of lineages are merged backwards in time with coalescence times drawn from 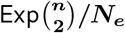, where ***n*** is the current number of lineages and ***N***_***e***_ is the effective population size. A stem branch ***t***_stem_ ~ Exp**(*N***_***e***_) is assigned to the root, representing the time from tumor founding to the most recent common ancestor. This allows clonal mutations to be included in the simulated dataset. ***M*** somatic mutations are placed on branches proportional to branch length, but because coalescent trees concentrate most branch length near the leaves, an optional uniform-CCF reweighting spreads mutations across the full CCF spectrum: eligible branches are grouped into ***K*** equal-width CCF bins and the sampling weight of branch ***i*** in bin ***b*** becomes

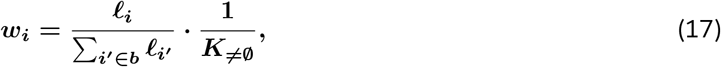

where **𝓁**_***i***_ is the branch length and ***K*** _**≠ ∅**_ is the number of non-empty bins. This preserves relative branch-length weighting within each CCF stratum while equalizing total probability across strata. Copy number ***C***_***j***_ ∈ **{1, 2, 3}** at each locus is drawn from a categorical distribution and multiplicity is sampled from a geometric prior 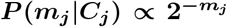, ***m***_***j***_ ∈ **{1, …**, ***C***_***j***_**}**, consistent with the inference model. Branches yielding CCF ***<* 0.05** are redrawn to avoid degenerate cases where sparse single-cell coverage is uninformative. Additionally, for each mutation, bulk reads are drawn from a beta-binomial with the expected VAF and single-cell reads are drawn depending on the chosen error model. With the beta-binomial model, a locus-specific mean concentration ***µ***_***τ***_ is drawn uniformly from **[*µ***_***τ***,**min**_, ***µ***_***τ***,**max**_] for each mutation to reflect genomic variation in amplification efficiency, and each cell’s concentration is then drawn as ***τ***_***i***_ ~ Gamma**(*µ***_***τ***_, CV_***τ***_ ). This per-cell ***τ***_***i***_ determines overdispersion for both germline SNPs and somatic variants, ensuring that bias estimated from hSNPs transfers to variant calls.

With the Lodato model, single-cell reads are generated in two steps. First, the Lodato amplification model produces a distorted read count at the mapped allele frequency ***θ***_***s***_ (equation (7)). Then, per-read sequencing error at rate ***ϵ***_***sc***_ is applied, matching the generative process of equation (8).

Table 1 shows the parameter values used for simulating data and performing inference. Copy number of each mutation was allowed to vary to test the severity of violating the diploid assumption of ProSolo. Coverage values for single-cells and the bulk were chosen to mimic values from typical empirical datasets. The sequencing error rate was set to 0.01, which is slightly higher than expected for a typical Illumina dataset [14]. This was chosen to account for the potentially increased sequencing error in single-cell experiments.

**Table 1:**
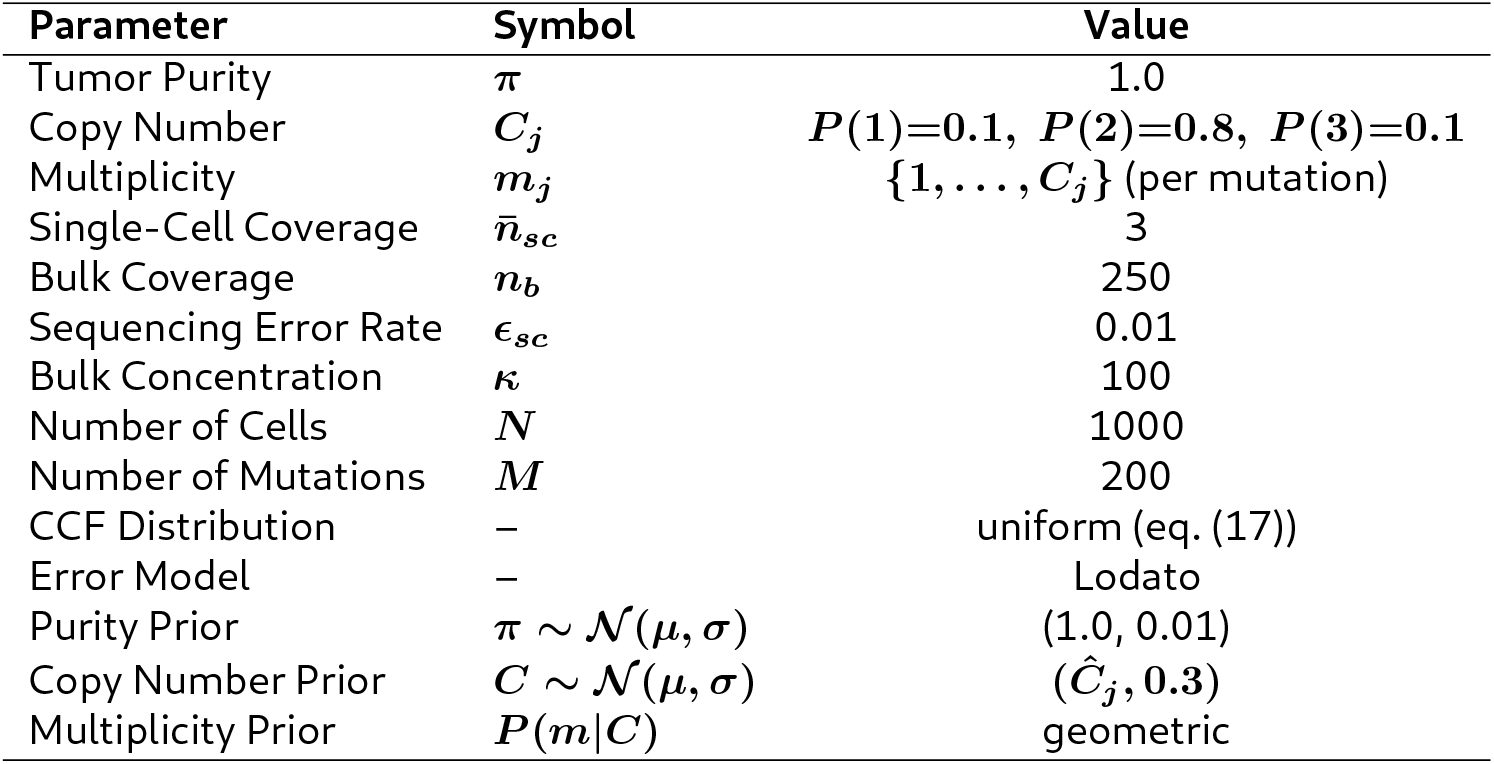
Simulation and inference parameters. Copy number and multiplicity vary per mutation and the copy number prior mean *Ĉ*_*j*_ is set to each mutation’s true value.

### 2.3 Implementation & Evaluation

SC-BIG is available as an open-source package at https://github.com/ICCB-Cologne/sc-big under the GNU General Public License v3. The model is implemented as a Python package providing both a library API and command-line interface with subcommands for simulation (`simulate`, `simulate-tree`), inference (`infer`, `infer-prosolo`), and method comparison (`compare`). All simulation hyperparameters and inference settings are configurable via command-line arguments.

Bayesian inference uses Pyro [15], a probabilistic programming library built on PyTorch. Pyro enables direct specification of the hierarchical model structure and provides a NUTS (No-U-Turn Sampler) implementation [16] for sampling of continuous parameters. Discrete parameters are enumerated and each MCMC sample receives a weight proportional to ***P* (*C*) · *P* (*m*** | ***C*) · *P* (*k***_***b***_ | ***n***_***b***_, CCF, ***π, C, m*)**. The hierarchical marginalization is computed as a weighted average over posterior samples. Single-cell inference is parallelizable across cells since each posterior probability is computed independently given bulk data, enabling efficient multi-core execution.

For comparison with ProSolo, the `infer-prosolo` subcommand generates synthetic BAM files from the simulated read counts using `pysam` and invokes the ProSolo v0.6.1 binary as a subprocess. For each cell at each mutation site, a minimal reference FASTA, candidate VCF, bulk BAM, and single-cell BAM data are generated with base qualities set to the Phred score corresponding to the simulated error rate (***Q* = – 10 log**_**10**_**(*e*)**). Importantly, when the Lodato error model is selected, the simulation generates single-cell reads from the same two-step process that ProSolo assumes internally. The seven PHRED-scaled event posteriors emitted by ProSolo are converted to linear probabilities, and ***P* (**alt present) is computed as the union over the five alt-present events (HET, HOM_ALT, ADO_TO_REF, ADO_TO_ALT, ERR_REF), follow-ing equation 9 of Lähnemann *et al*. [9].

We evaluate classification performance using the complementary metrics. ROC-AUC (area under the receiver operating characteristic curve) is defined as follows:

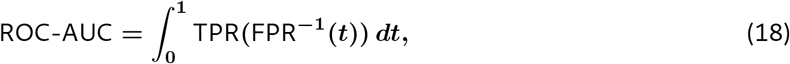

where TPR (true positive rate) **=** TP***/*(**TP **+** FN) and FPR (false positive rate) **=** FP***/*(**FP **+** TN). Precision-recall AUC (PR-AUC) is defined analogously as the area under the precision–recall curve, where precision **=** TP***/*(**TP **+** FP) and recall **=** TPR. For brevity, only the AUC metrics are presented here, but both metrics yield qualitatively similar results.

## 3 Results

We evaluated SC-BIG on simulated datasets with parameters defined in Table 1. Both procedures use the Lodato error model and for method comparisons, we present SC-BIG, ProSolo, and a naive caller that scores each cell by its observed VAF (***k***_***sc***_***/n***_***sc***_). All methods are evaluated using ROC-AUC and PR-AUC.

Figure 8 in the Appendix shows the simulated tree as well as the resulting mutation landscape. The CCF histogram (Figure 8b) confirms that the reweighting procedure (equation (17)) produces mutations across the full prevalence range. Copy number and multiplicity assignments (Figure 8c) follow their categorical and geometric priors, respectively, with the majority of mutations at ***C* = 2, *m* = 1**. The clustered heatmap (Figure 8d) reveals the expected clonal architecture. Groups of cells sharing nested sets of mutations, consistent with the underlying tree topology.

Given this simulated data, Figure 3 compares inference results from SC-BIG, ProSolo, and naive VAF thresholding across all 200 mutation sites. SC-BIG achieves the highest ROC-AUC of 0.942, followed by ProSolo (0.877) and naive VAF (0.869) (Figure 3, left panel). Calibration analysis (Figure 3, right panel) shows that SC-BIG’s posteriors closely track the ideal diagonal (slope **= 1.03, *R***^**2**^ **= 0.96**), while ProSolo exhibits both attenuation and scatter (slope **= 0.78, *R***^**2**^ **= 0.77**). Naive VAF shows intermediate performance with reasonable linearity (***R***^**2**^ **= 0.88**) and a large intercept (0.38).

**Figure 3:**
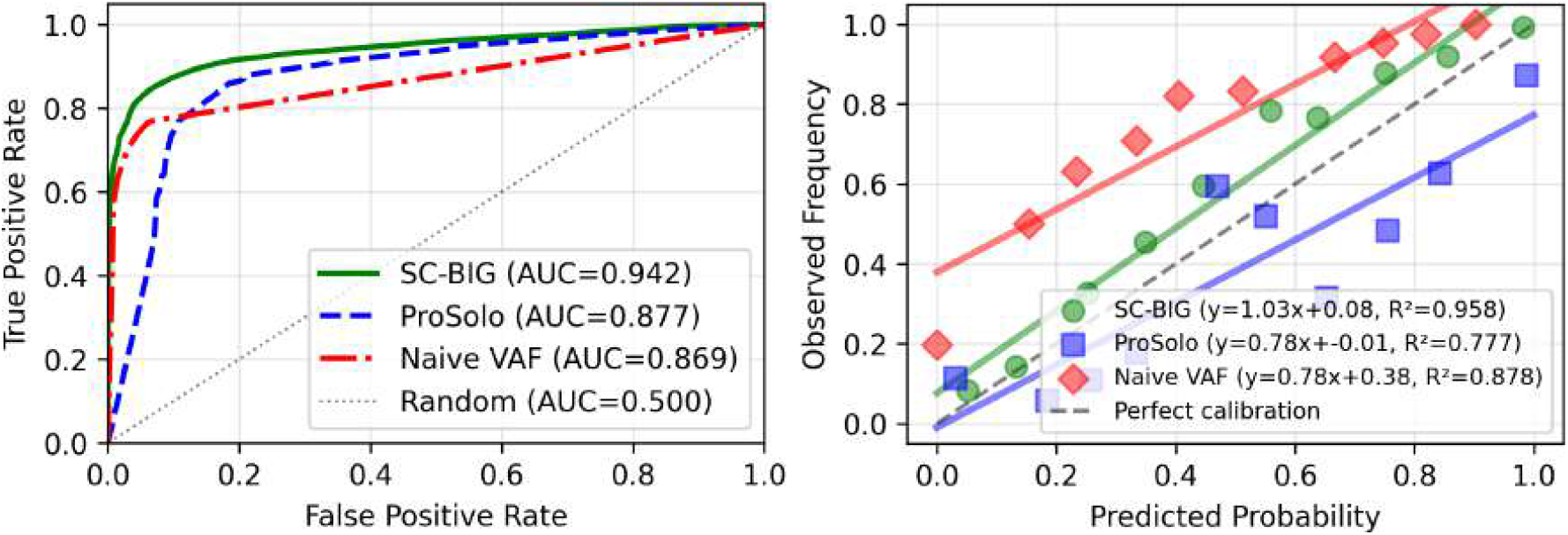
Overall method comparison. Left: ROC curves for SC-BIG, ProSolo, and naive VAF thresholding across all mutations. Right: Calibration plots with linear fits.

To investigate model performance with respect to cellular prevalence of the candidate mutation, we performed subset analysis as shown in Figure 4. For CCF ∈ **(0, 0.3]** (Figure 4, top-left), all three methods achieve comparable ROC-AUC with almost perfect calibration for SC-BIG. For CCF ∈ **(0.3, 0.7]** (Figure 4, top-middle), SC-BIGs ROC-AUC clearly separates from ProSolo and naive VAF (0.935 versus 0.693 versus 0.884). Notably, ProSolo’s ROC curves exhibit a sharp collapse at low false positive rates and low recall, respectively. A similar, yet less pronounced result emerges for CCF ∈ **(0.7, 1.0]** (Figure 4, top-right). ProSolo’s ROC-AUC drops to 0.700, below naive VAF (0.870) and SC-BIG (0.921). With respect to calibration (Figure 4, bottom row), SC-BIG appears to be well calibrated for medium and high CCF mutations, too.

**Figure 4:**
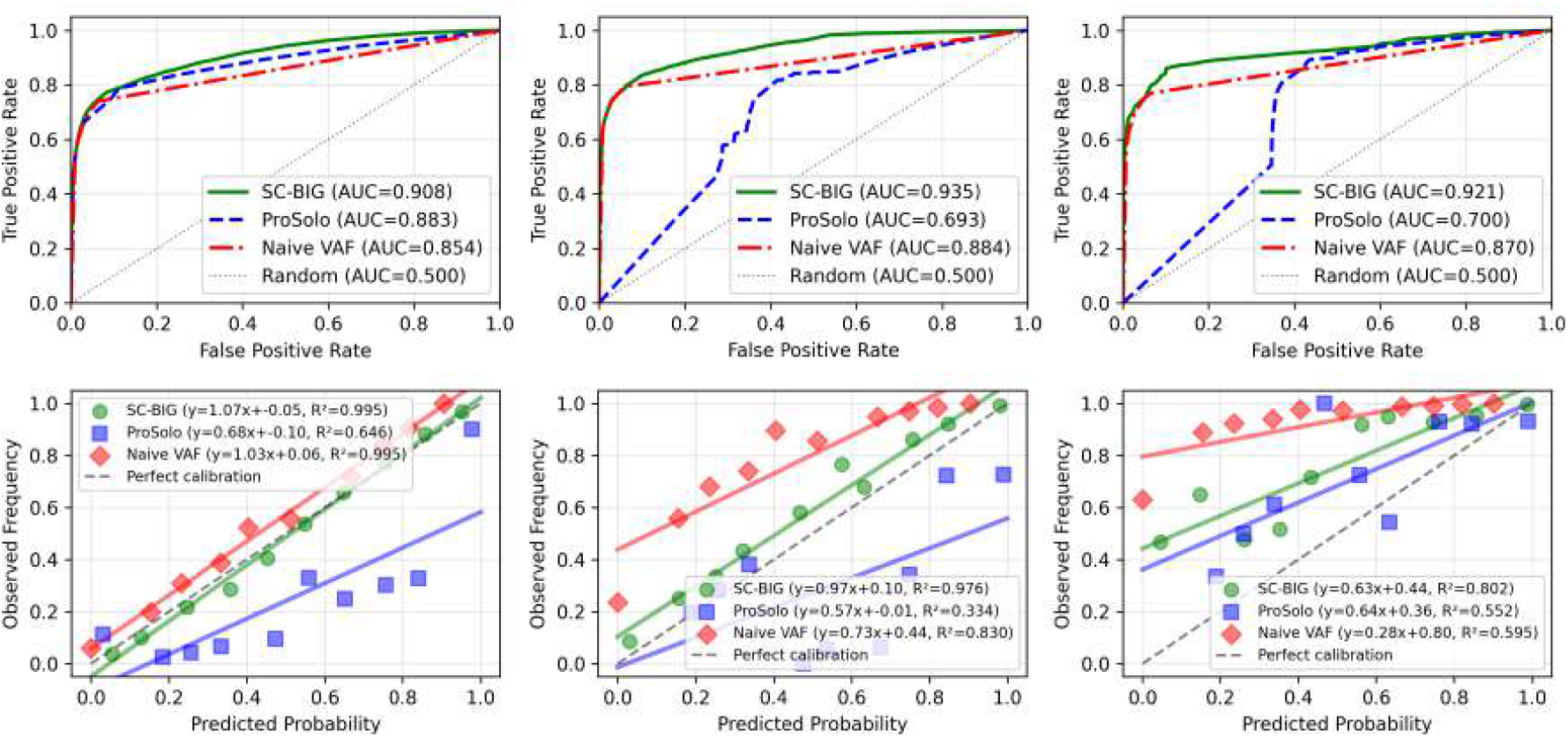
CCF-stratified method comparison. Left column: low CCF stratum (CCF ∈ **(0, 0.3]**). Middle column: mid CCF stratum (CCF ∈ **(0.3, 0.7]**). Right column: high CCF stratum (CCF ∈ **(0.7, 1.0]**). Rows show ROC curves (top) and calibration plots (bottom).

Whereas this first simulation of a pure bulk tumor represented ideal bulk evidence, we performed a similar experiment with ***π* = 0.5** as well. While SC-BIG still outperforms ProSolo in this scenario, ROC-AUC differences are less marked (Figure 5). Additionally, ProSolo does not show performance degradation for medium and high CCF ranges (Figure 6).

**Figure 5:**
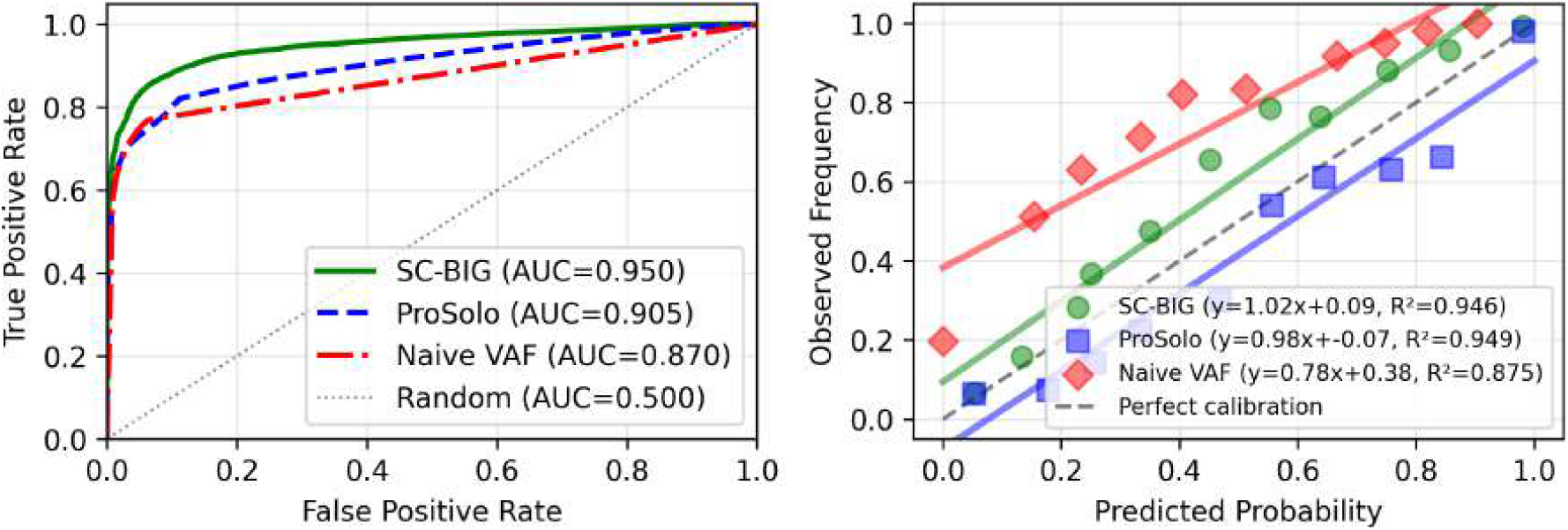
Overall method comparison for an impure tumor (*π* = 0.5). Left: ROC curves for SC-BIG, ProSolo, and naive VAF thresholding across all mutations. Right: Calibration plots with linear fits.

**Figure 6:**
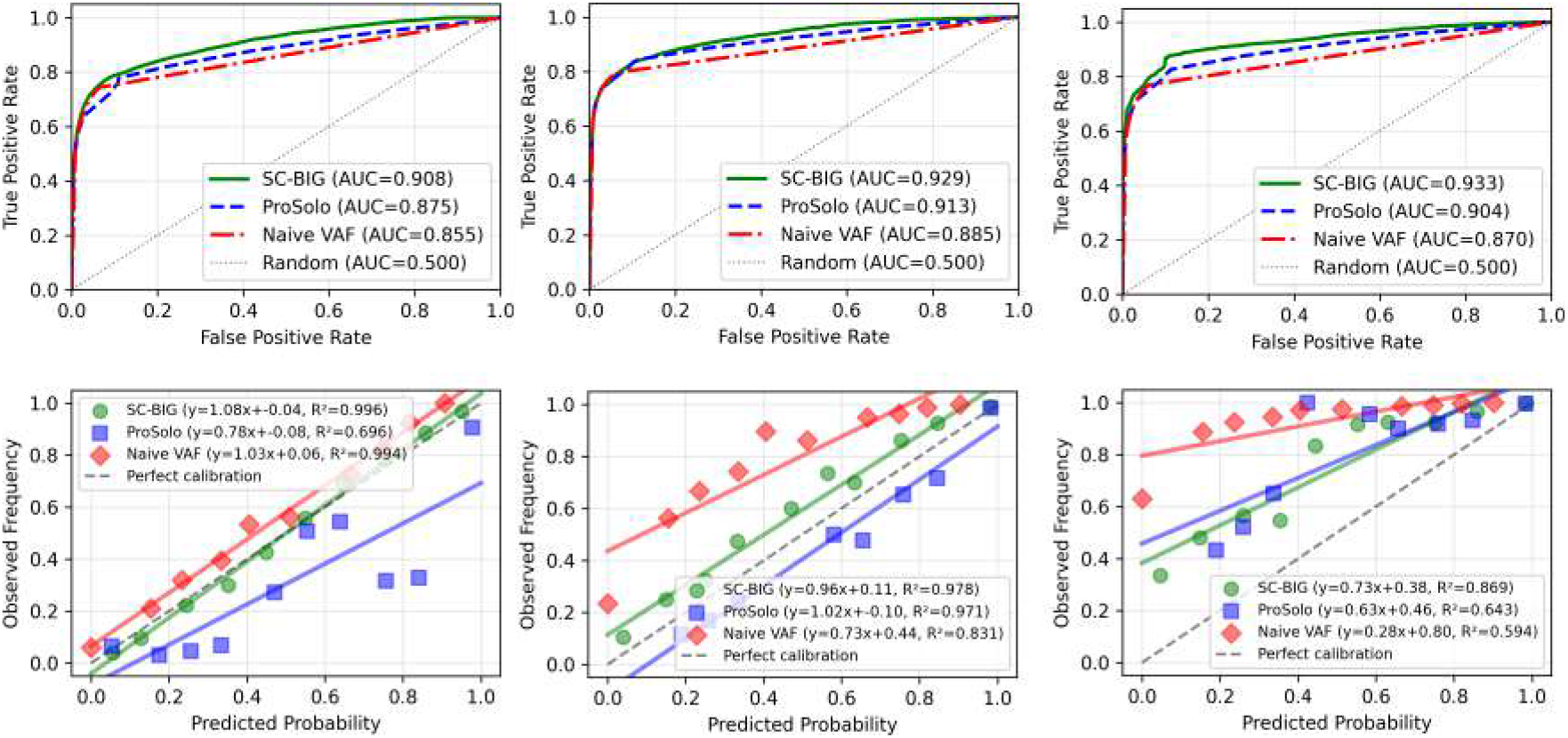
CCF-stratified method comparison for an impure tumor (*π* = 0.5). Left column: low CCF stratum (CCF ∈ **(0, 0.3]**). Middle column: mid CCF stratum (CCF ∈ **(0.3, 0.7]**). Right column: high CCF stratum (CCF ∈ **(0.7, 1.0]**). Rows show ROC curves (top) and calibration plots (bottom).

## 4 Discussion

Here, we present SC-BIG, a Bayesian model for bulk-informed single-cell variant calling. We provide a simulation framework to test SC-BIG and compare its performance ProSolo and with naive VAF thresholding. In these comparisons, SC-BIG outperforms both methods across the entire CCF spectrum. Importantly, SC-BIG and ProSolo use the same Lodato per-read error model [11] so that differences in discrimination reflect modeling assumptions rather than error-model mismatch. In addition to improved classification metrics, SC-BIG provides interpretable uncertainty quantification through well-calibrated posterior probabilities. It outputs a matrix of genotype probabilities that quantifies confidence in variant presence, and calibration analysis demonstrates that a cell assigned posterior probability ***p*** truly harbors the variant with frequency **≈ *p*** across all tested CCF regimes. This calibration facilitates threshold selection based on the tolerance of a specific experiment for false positives versus false negatives. Additionally, calibrated posteriors enable integration into downstream probabilistic analyses. Phylogenetic reconstruction methods such as CellPhy [17] can incorporate genotyping uncertainty via genotype likelihoods as input, yielding more accurate trees than methods that require binary genotype calls.

In our simulations, ProSolo performance collapsed for medium and high CCF variants. We hypothesize that its restricted genotype mixture model favors near-clonal variant interpretations, degrading performance as bulk CCF rises. At low CCF with high tumor purity, the weak bulk signal causes ProSolo to appropriately assign low variant probabilities, achieving ROC-AUC close to SC-BIG. At mid-(**0.3 – 0.7**) and high-range (**0.7 – 1.0**) CCF on the other hand, the model begins favoring explanations in which most tumor cells carry the variant, likely explaining non-carrier cells as ADO rather than true negatives. This behavior likely arises because ProSolo’s bulk model restricts allele frequencies to mixtures of adjacent diploid genotypes, biasing inference toward widespread variant presence once the bulk signal is strong. This results in bulk information actively degrading performance compared to ignoring it entirely. Notably, at reduced tumor purity (***π* = 0.5**), the weaker bulk signal attenuates the miscalibration effects described above, thus supporting our hypothesis.

SC-BIG instead marginalizes over CCF as a continuous latent variable, recognizing that even at high prevalence, a fraction of cells may not carry the variant. The performance gap is potentially further widened by copy number and multiplicity uncertainty. The observed bulk VAF is consistent with multiple **(*C, m*)** configurations, and SC-BIG’s joint marginalization over these discrete states can yield more accurate per-cell posteriors than ProSolo’s fixed diploid model.

The presented work is limited in several ways. First, the comparison with ProSolo, while controlled, involves design choices that favor SC-BIG. The copy number and purity priors are set to each mutation’s true values with little variance, providing SC-BIG with accurate information that might not be available in practice, because estimates for ***C*** and ***π*** derived from real-world data can carry substantial uncertainty. Additionally, ProSolo is invoked on synthetic BAM files generated from simulated read counts. While we have verified that this pipeline produces valid alignments, subtle errors could in principle affect ProSolo’s performance. The same holds true for the Lodato error model which we have implemented carefully from its description in the respective publications but which might contain undetected software bugs.

Second, the empirical CCF prior (equation (12)) is estimated from the same single-cell data that are subsequently genotyped. While the prior summarizes population-level prevalence and does not use per-cell genotype information, it introduces a certain degree of circularity. We conjecture that this effect will be small in practice, but future research needs to address this limitation.

Third, the Lodato error model maps the true allele frequency ***m/C*** to one of three discrete values **{0, 0.5, 1}** (equation (7)). This discretization loses information for non-diploid states. While SC-BIG’s technology-agnostic beta-binomial model (equation (6)) does not suffer from this limitation, it is unlikely to fit data well in a non-simulation context. This is especially true because it assumes a 1:1 allelic ratio which might not hold for loci affected by copy number changes and because we do not investigate genomic windows suitable for the estimation procedure. In the future, a method akin to the one described in Luquette *et al*. [4] could be used to generalize SC-BIG’s error models to non-MDA technologies.

Further developments could include incorporation of per-cell copy number estimates as informative priors on ***C***, reducing dependence on bulk-derived estimates and improving inference for aneuploid tumors where copy number varies across cells. Phylogenetic constraints—for example, the requirement that mutations in a subclone also appear in ancestral cells—would most likely further improve genotyping accuracy. Testing of SC-BIG on empirical datasets will allow us to identify how applicable the model is to real experimental scenarios. Real data validation is essential as it introduces systematic biases and model violations that simulations cannot capture.

## 5 Acknowledgements

DS is partially funded through a research grant by the E.I. Stiftung Kölner Krebsforschung. RFS is a Professor at the Cancer Research Center Cologne Essen (CCCE) funded by the Ministry of Culture and Science of the State of North Rhine-Westphalia. This work was supported by the Bruno and Helene Jöster Foundation as “CLONETRAC - Tracking the clonal dynamics of cancer through treatment at the single-cell level”. The authors thank the Regional Computing Center of the University of Cologne (RRZK) for providing computing time on the High Performance Computing (HPC) system RAMSES as well as support. The large language model Claude Sonnet 3.5 was used for proof-reading and improving the clarity of the manuscript [18]. The SC-BIG logo was created with the help of GPT-5 mini [19].

## 7 Appendix

SC-BIG takes as an input bulk and single-cell sequencing data, derived bulk data like copy number and purity estimates, and error parameters. Figure 7 demonstrates the relationship between the directly observed parameters.

**Figure 7:**
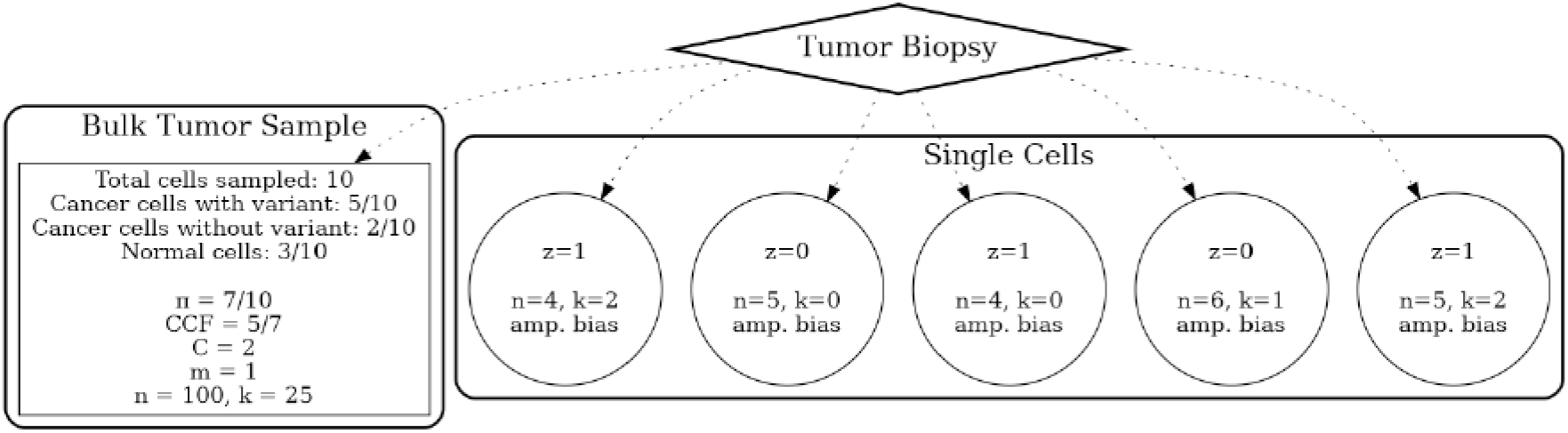
Biological interpretation of model parameters. The bulk tumor contains a mixture of cancer and normal cells, with only a fraction of cancer cells harboring the variant (CCF). Copy number state ***C***, purity ***π***, and variant read counts ***k*** are sampled with uncertainty. Germline SNPs **ℋ** provide amplification bias estimate under one of SC-BIG’s error models.

Bayesian inference of variant presence then proceeds according to algorithm 1.

### Algorithm 1

Hierarchical Bayesian Inference

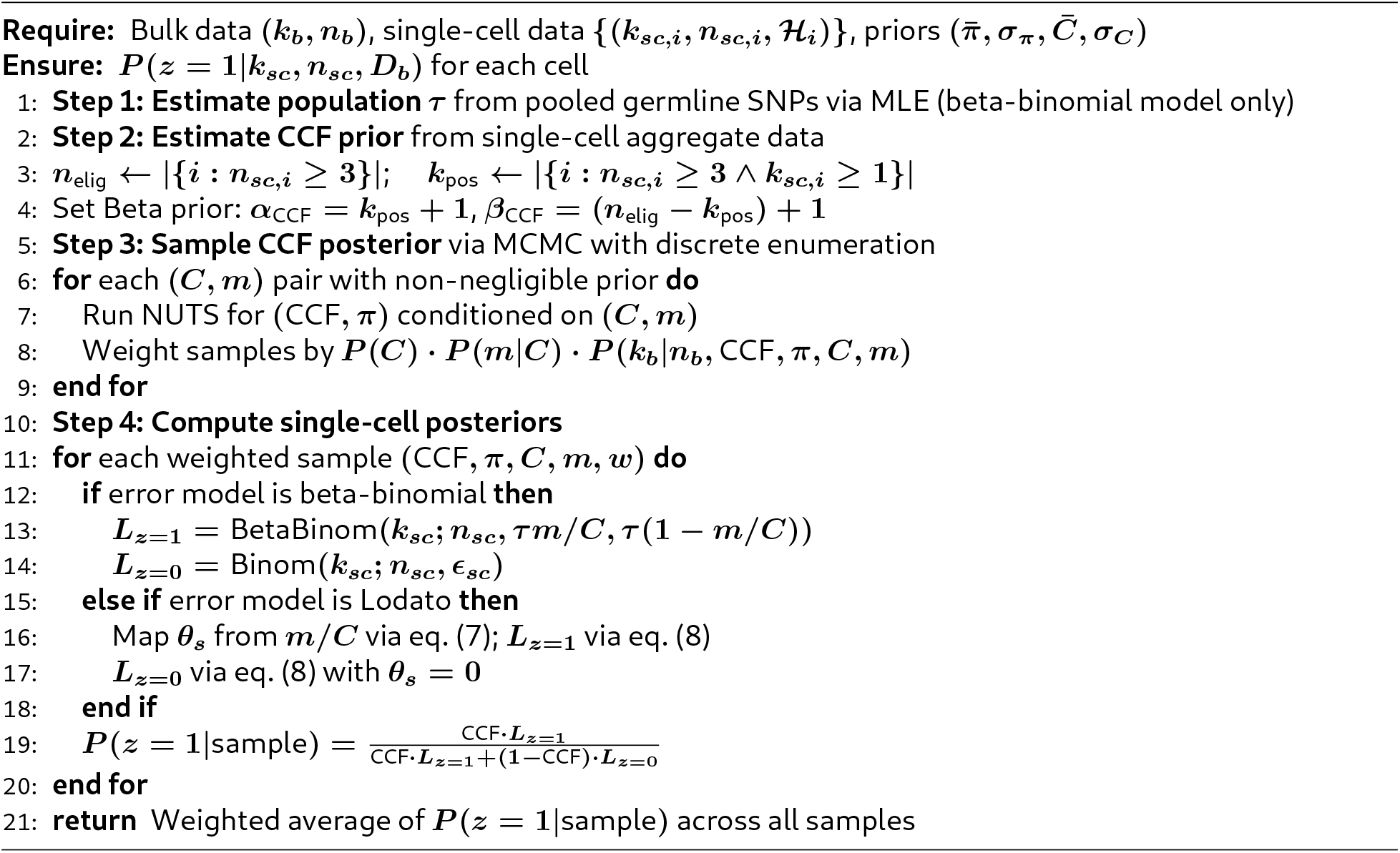

To test SC-BIG, a simulation procedure is used that generates a population of cells harboring mutations with different levels of subclonality. Algorithm 2 gives a detailed description of the employed process.

Figure 8 summarizes key characteristics of the cell population produced by the hyperparameters of Table 1.

**Figure 8:**
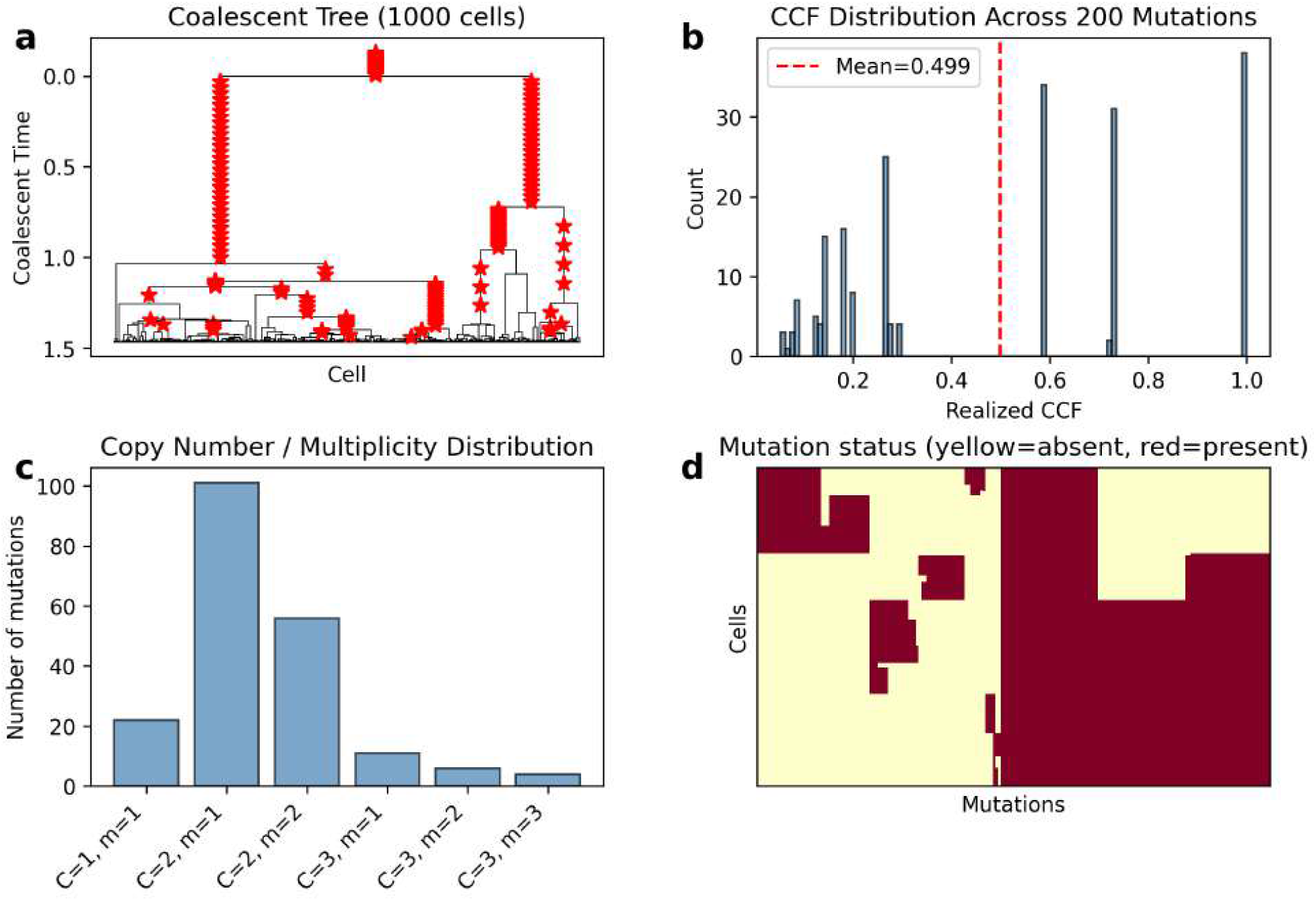
Coalescent population simulation. (a) Coalescent tree with ***N* = 1000** cells; red stars mark mutations. (b) Distribution of realized CCF values across ***M* = 200** mutations. (c) Copy number and multiplicity assignments. (d) Clustered cells **×** mutations heatmap showing variant presence (red) and absence (yellow).

### Algorithm 2

Coalescent Simulation

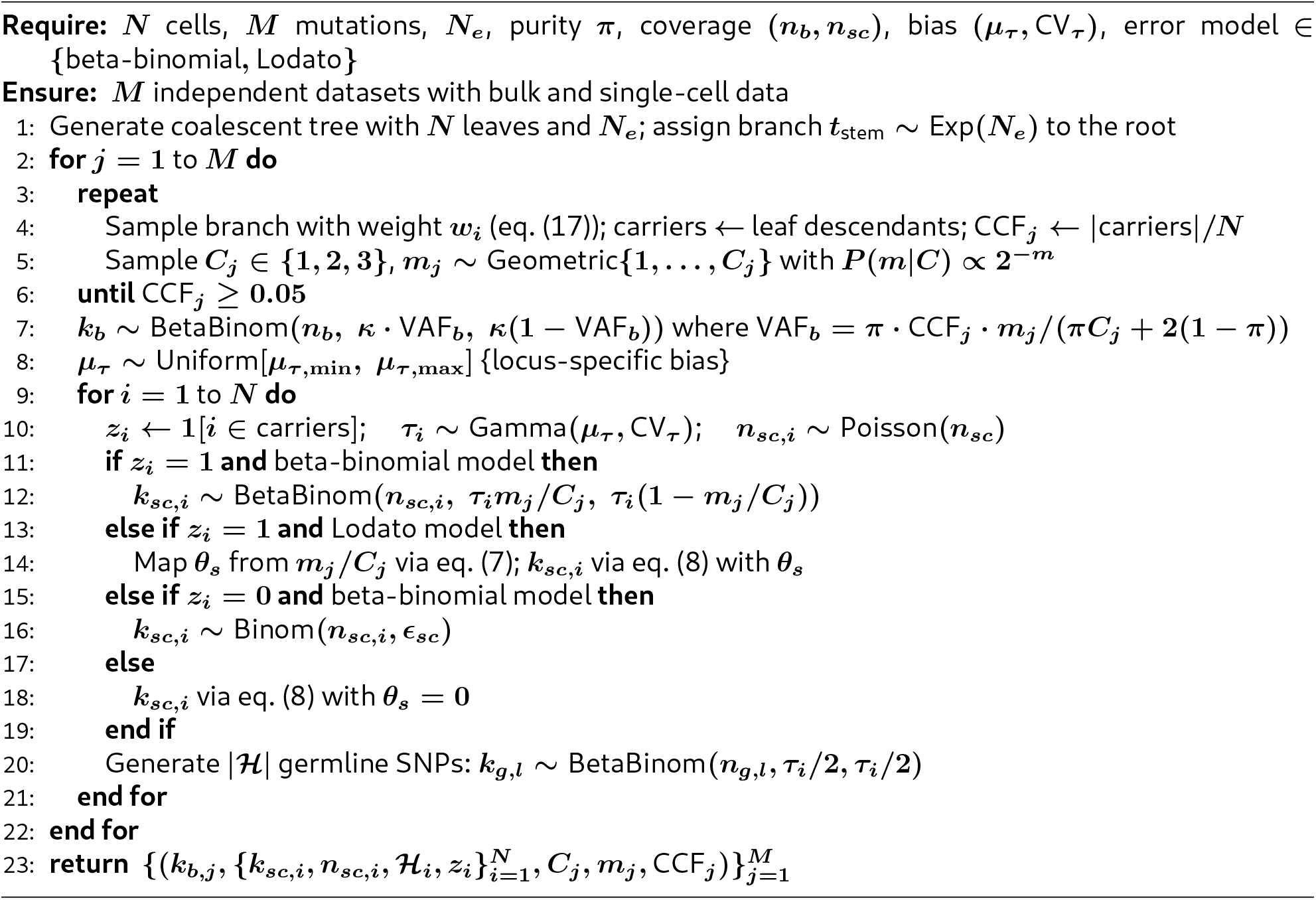

## Bibliography

1. Wu, F. et al. Single-cell profiling of tumor heterogeneity and the microenvironment in advanced non-small cell lung cancer. Nature Communications 12, 2540. ISSN: 2041-1723 (May 2021).

2. Zafar, H., Wang, Y., Nakhleh, L., Navin, N. & Chen, K. Monovar: single-nucleotide variant detection in single cells. Nat. Methods 13, 505–507 (June 2016).

3. Dong, X. et al. Accurate identification of single-nucleotide variants in whole-genome-amplified single cells. Nat. Methods 14, 491–493 (May 2017).

4. Luquette, L. J., Bohrson, C. L., Sherman, M. A. & Park, P. J. Identification of somatic mutations in single cell DNA-seq using a spatial model of allelic imbalance. Nat. Commun. 10, 3908 (Aug. 2019).

5. Rozenblatt-Rosen, O. et al. The Human Tumor Atlas Network: Charting tumor transitions across space and time at single-cell resolution. Cell 181, 236–249 (Apr. 2020).

6. Leighton, J., Hu, M., Sei, E., Meric-Bernstam, F. & Navin, N. E. Reconstructing mutational lineages in breast cancer by multi-patient-targeted single-cell DNA sequencing. Cell Genom. 3, 100215 (Jan. 2023).

7. Chen, W. et al. Analysis of genomic heterogeneity and the mutational landscape in cutaneous squamous cell carcinoma through multi-patient-targeted single-cell DNA sequencing. BMC Cancer 25, 1362 (Aug. 2025).

8. Zhang, H. et al. Application of high-throughput single-nucleus DNA sequencing in pancreatic cancer. Nat. Commun. 14, 749 (Feb. 2023).

9. Lähnemann, D. et al. Accurate and scalable variant calling from single cell DNA sequencing data with ProSolo. Nature Communications 12, 6744. ISSN: 2041-1723 (Nov. 2021).

10. Köster, J., Dijkstra, L. J., Marschall, T. & Schönhuth, A. Varlociraptor: enhancing sensitivity and controlling false discovery rate in somatic indel discovery. Genome Biology 21, 98. ISSN: 1474-760X (Apr. 2020).

11. Lodato, M. A. et al. Somatic mutation in single human neurons tracks developmental and transcriptional history. Science 350, 94–98 (Oct. 2015).

12. Tarabichi, M. et al. A practical guide to cancer subclonal reconstruction from DNA sequencing. Nature methods 18, 144–155. ISSN: 1548-7091 (Feb. 2021).

13. Kingman, J. F. C. The coalescent. Stoch. Process. Their Appl. 13, 235–248 (Sept. 1982).

14. Stoler, N. & Nekrutenko, A. Sequencing error profiles of Illumina sequencing instruments. NAR Genomics and Bioinformatics 3, lqab019. ISSN: 2631-9268 (Mar. 2021).

15. Bingham, E. et al. Pyro: Deep Universal Probabilistic Programming. J. Mach. Learn. Res. 20, 28:1–28:6 (2019).

16. Hoffman, M. D. & Gelman, A. The No-U-Turn Sampler: Adaptively Setting Path Lengths in Hamiltonian Monte Carlo (Nov. 2011).

17. Kozlov, A., Alves, J. M., Stamatakis, A. & Posada, D. CellPhy: accurate and fast probabilistic inference of single-cell phylogenies from scDNA-seq data. Genome Biology 23, 37 (2022).

18. Anthropic. Introducing the next generation of Claude 2024. https://www.anthropic.com/news/claude-3-family.

19. OpenAI. ChatGPT (GPT-5 Mini) 2026. https://openai.com/chatgpt.

